# Disparities in information on Long-Acting Reversible Contraceptives available to college students on student health center websites in USA

**DOI:** 10.1101/2020.01.27.920926

**Authors:** Anagha Kulkarni, Tejasvi Belsare, Risha Shah, Diana Yu Yu, Carrie Holschuh, Venoo Kakar, Sepideh Modrek, Anastasia Smirnova

## Abstract

Long Acting Reversible Contraceptive (LARC) methods are among the most effective birth control approaches for adolescent and young adults yet information on these methods is not widespread. We examine LARC information provided by Student Health Centers (SHC) websites from Universities across the USA to document disparities in access to information on these important contraception methods for college students. We find that compared to EC, Condoms, (plus Pap smear as control), LARC is mentioned less frequently than the others and 73% of schools have no LARC content on their SHC websites. There is no standardization in how the sexual and reproductive health information is organized on SHC websites, which might hinder access. When LARC information does exist, readability and accessibility vary. Universities having high rates of the student body who are African American or female are less likely to provide LARC information on their SHC website and universities situated in more rural settings are less likely to post LARC information on their websites.

## 1 Introduction

Universities have historically been reluctant partners in providing access to reproductive services through Student Health Centers (SHCs). In the U.S., external social and political forces have played a major role in shaping SHCs’ provision of contraception care. Student activists have been the driving force pushing universities and their health centers into providing these important services since the 1970s [1]. Laws like Title IX, the 1972 civil rights law prohibiting sex discrimination in federally funded educational programs, have been used to argue that SHCs must provide access to contraception in order to maintain equal access to education based on sex. In addition, the enactment of the Affordable Care Act (ACA) in 2010 required coverage of contraception as an essential health benefit, thereby changing the contraception care landscape in the U.S., including at universities. This led to schools either dropping student insurance plans altogether (such as Brigham Young University) or complying with the mandate of providing contraception coverage. The ACA prioritized reproductive health care in an effort to reduce existing socioeconomic barriers to essential preventive care for women, in the hope of thereby addressing associated health-related disparities [2]. There is now evidence that impacts of the ACA include narrowing the gap in prescription contraception access between black and white women, where black women have historically accessed prescription forms of contraception at lower rates than white women [3].

SHCs must both provide services and adequately inform the student population about these services in order to ensure equitable access to reproductive health care for college students [4]. While much progress has been made in increasing basic access to contraception thorough SHC clinics and provider appointments [5, 6], new barriers are emerging. Rather than physically visiting an SHC location on a university campus, most students now make their first contact with an SHC through the internet [7, 8].

### 1.1 Student health centers as sources of health information

SHCs and their websites are well positioned to serve as an equitable source of high-quality information to adolescent and young adult students [9]. As of 2016, approximate 41% of 18 to 24-year-olds were enrolled in college with a higher proportion of female than male attendees (43% female vs. 38% men) and growing racial/ethnic diversity of the student population. Moreover, in studies of sources of health information amongst college students, black and Hispanic students were more likely than white students to use SHC staff as a source of health information [10]. This suggests that as the student bodies of universities diversify, SHCs may take on a larger role in supporting student health through the equitable provision of appropriate education and services.

Given the central role that SHCs play in providing health information and services to adolescents and young adults, there is growing interest in documenting the services and information that they provide. A recent study that surveyed college SHCs about their provision of sexual health services had a response rate of about 55% [6], reflecting a strong level of interest in the issue of providing sexual health services through SHCs. However, a key limitation in survey-based studies is that response rates may be biased by issues such as the role of the survey respondent, the level of support for contraceptive care at the SHC, or the resources devoted to reproductive health services at a particular campus. To assess SHCs’ role in contraceptive access and education for students, an alternative strategy to administering a survey would be to systematically assess information that SHCs provide to students through their websites. This approach encompasses the online nature of most students’ searches for contraceptive information and services. It may allow for the assessment of factors which impact students’ ability to access high-quality information about effective and appropriate contraceptive options.

### 1.2 Long-acting reversible contraception for the college-aged population

Long-acting reversible contraceptive (LARC) methods are among the most effective birth control approaches [11]. LARC contraceptive methods include intrauterine devices (IUD), the hormonal implant, and the shot. Nationally their use as primary source of contraception has been rapidly increasing from 2.4% in 2002 to 11.6% in 2012, a 5-fold increase in 10 years [12, 13]. Updated guidelines recommend these methods for use in adolescents and young adults, as they are considered to be extremely effective and safe for this population at high risk of unintended pregnancy [11]. The increase in use has been most rapid for low-income adolescents, increasing 18-fold from 2005-2015 in 15-19 year old women who use federally funded clinics [14]. Meanwhile the rate of unintended pregnancy in the United States has declined substantially. In particular, teen birth rates have fallen by 53% between 2007 and 2015 [15]. Nobles et al. reported 15% higher Google searches for IUDs from November 1, 2016 to October 31, 2017 than what was expected based on prior years’ search trends, indicating increased awareness and interest in LARC methods among the overall population of the country [16].

The question remains as to whether the increase in LARC use has contributed to the decline in teen birth rates. In spite of the last decade’s increase in LARC usage, overall LARC adoption rates remain relatively low, especially among adolescents. Less than 6% of U.S. adolescents have used LARC methods [17, 18]. This is even more perplexing when the following factors are considered: 1. Adolescents and young adults are at a high risk of unintended pregnancy [15, 19, 20], 2. The U.S. continues to have the highest adolescent birth rates of any developed country [20], and 3. Unintended pregnancy has extremely severe lifetime consequences for adolescents, including the interruption or prevention of meaningful education [15, 19, 20].

Accurate information and adequate support in decision-making have been shown to significantly influence women’s successful use of contraception [8, 21]. Clinician bias or misinformation can be a barrier to adequate LARC access for adolescents and young adults [22, 23]. Lack of knowledge was identified as the most common barrier to LARC use in one survey of 1,982 female undergraduate students. LARC usage among college students, while increasing, continues to trail usage of older and less reliable methods [4, 24]. Many college students report concern over unplanned pregnancy as well as a lack of sufficient LARC education and access [4, 25], as well as a desire for greater access to more effective methods including LARC [26]. Misconceptions about infertility and other harms have been identified as a common barrier to IUD uptake among female college students [27]. SHCs and their websites are well positioned to provide high-quality LARC information to diverse populations of students [9].

In this study, we take an equity lens and systematically document the information provided by SHC websites at 4-year public universities across United States. We focus on public institutions as they are obligated under Title IX to limit sex-related barriers and required under ACA to provide coverage to contraception if they offer a student health plan. We undertake 3 distinct studies/analyses that build on one another to examine access to online information about LARC. In our first study we examine information on SHC websites on LARC relative to other forms of contraception and reproductive health services. In our second study we develop and deploy an algorithm to search for LARC information on SHC website in a systematic way. In a third study, we examine how the online information for LARC varies by university-level characteristics.

## 2 Study 1: Comparing information on common reproductive health services on SHC websites

The goal of this study is to understand the prevalence of LARC information on SHC websites and to compare it the prevalence of information of other contraception methods and common reproductive health services on SHC websites.

### 2.1 Study 1: Methods

For this study we focused on four categories: LARC methods, Emergency Contraception (EC) methods, Condoms, and Pap smear. Information about EC and condoms was assessed in order to provide a comparison between LARC and less effective, but more commonly used contraceptive methods. Pap smear information was measured in order to compare SHCs’ contraceptive information with their provision of information regarding a routine, preventive, non-contraceptive reproductive health service.

Current literature regarding evidence-based standards for LARC in the U.S. [11] was used to determine a list of available LARC methods (Intrauterine devices (hormonal and copper), implant, and shot) and a range of acceptable terms for each method, including brand names, pharmaceutical language and popular terms. This led to the development of the following LARC keyword list: *IUD, Progesterone IUD, Progestin, Hormonal IUD, Mirena, Skyla, Kyleena, Liletta, Copper IUD, Non-Hormonal IUD, Paragard, (contraceptive) implant, Nexplanon, (contraceptive) injection, Shot, Depo-Provera OR Depo*. Similarly for EC methods, the following keyword list was developed: *emergency contraception/contraceptives, morning after pill, Plan B (levonorgestrel), ella (ulipristal acetate), copper IUD, Paragard, non-hormonal IUD*. The keyword list for condoms simply consisted of the term *condom*. Similarly for Pap smear it was *pap smear*.

The set of educational institutions for this study were chosen by first querying the National Center for Educational Statistics (https://nces.ed.gov/collegenavigator/) to identify all 4-year, public institutions that grant bachelor’s degrees in the U.S. This retrieved a set of 591 universities. Next, a subset of 200 universities was randomly sampled from the 591 universities to define the final set for this study (File S1 File.).

The SHC websites of the selected 200 universities were then analyzed manually for presence of the four categories under study (LARC, EC, Condoms, Pap smear). Specifically, annotators were instructed to: 1. locate the SHC website for a given university, and 2. then search for the specified category’s keywords on the SHC website. Presence of any of the keywords from the category was recorded as value *1* for that category, and absence of all keywords was recorded as value *0* for that category. For example, if a SHC website mentioned *Mirena* on one or more web-pages, then the annotator would report *1* for the LARC category. Every category was annotated by two independent coders. Cohen’s kappa coefficient was computed for each category to measure inter-annotator reliability.

For subsequent analysis we computed *Average Annotation Value (AAV)* for each data point (university) which is the average of the coders’ response value. For example, if both annotators agreed that one or more keywords were present for a category, then then average was 1, if they agreed that keywords were absent, the average was 0, and if they disagreed, the average was 0.5. The AAV essentially reflects the uncertainty coming from the disagreement, while allowing us to keep the data point. This leads to generation of 200 AAVs for each category. Using these AAVs, the LARC category were compared with each of the remaining three categories (EC, Condoms, Pap smear) for systematic differences using two-tailed paired t-test.

### 2.2 Study 1: Results

The results of the comparative analysis of the four categories are reported in Table 1. The average of the 200 AAVs for each category is reported under the column *Mean*. These values suggest that the overall prevalence of all four categories is low on SHC websites. The lowest prevalence is for LARC methods (0.29), closely followed by EC (0.30), Condoms (0.35), and Pap smear (0.49). The presence of Pap smear is significantly more prevalent that that of LARC methods on SHC websites (*p*<0.01). The difference between LARC and Condoms prevalence is marginally significant (*p*<0.1).

**Table 1.** Summary statistics and significance testing for difference in prevalence of information of LARC methods compared to information of Emergency Contraception (EC), Condoms, and Pap smear, on SHC website. The *p*-values come from two-tailed paired t-tests.

|  | Mean | SD | t | df | p |
| --- | --- | --- | --- | --- | --- |
| LARC | 0.29 | 0.41 | . | . | . |
| EC | 0.30 | 0.41 | 0.49 | 187 | 0.62 |
| Condoms | 0.35 | 0.41 | 1.78 | 187 | 0.08 |
| Pap smear | 0.49 | 0.44 | 6.27 | 187 | <0.01 |

If the three categories, EC, Condoms, and Pap smear, are collectively compared with LARC then the difference is statistically significant (t(187) = 3.25, *p* = 0.001, two-tailed) with LARC (M = 0.29, SD = 0.41) and collective category (M = 0.38, SD = 0.35).^1^ LARC methods were mentioned less frequently on average than the other methods.

The inter-annotator reliability results are given next. Cohen’s kappa coefficients were 0.64, 0.60, 0.44, and 0.51 for LARC, EC, Condoms, and Pap smear, respectively. These values are typically interpreted as moderate agreement. It could be argued that the coefficient for LARC (0.64) is bordering on substantial agreement.

We had expected much higher coefficients since we believed that the annotation task (searching for specified keywords on a website) was straightforward and objective. Neither of these assumptions were found to be true. Annotators reported that the placement of relevant information was not always intuitive, which leads to annotation errors. There is high variability in how the content on the SHC websites is organized, which makes the annotation task longer, tedious, and further amplifies the possibility to human error. The highest agreement on LARC methods also suggests that unconscious bias might have been introduced into the annotated data – annotators might have been extra careful when searching for LARC methods since it is the focus of the study. These trends motivate the next study which employs computational approaches to avoid human error and bias.

## 3 Study 2: Automated methods to gather LARC data on national sample of university SHC websites

The findings from Study 1 and our experience of conducting Study 1, motivates this next study where the central goal is to expand the LARC data gathering efforts in order to analyze national-level trends. Computational approaches lend well to this goal since the LARC data that needs to be gathered is available in digital format on the World Wide Wed (WWW), specifically the university student health center websites. Employing computational approaches offers two important benefits: 1. efficient scaling-up, and 2. effective tracking. Once the computational approaches are designed and developed they can be applied to as many universities as needed without any additional cost (efficient scaling-up). These approaches can also be re-applied as many times as needed to track changes in the data (effective tracking). Achieving either of this with traditional data gathering instruments such as, surveys or annotation efforts (Study 1) is extremely difficult, inefficient, and expensive.

In this study we have developed computational approaches for 1. identifying the student health center website for a given university, 2. assessing the accessibility of LARC information available on SHC websites, and 3. assessing the quality of the LARC information available on SHC websites.

### 3.1 Study 2: Methods

We commence the study by defining the set of universities that will be analyzed using the computational approaches. As in Study 1, we started with the list of all 4-year, public, bachelor’s granting institutions in the US. A list of 591 institutions. From this list, nine Native American serving institutions and Veteran serving institutions were dropped because they have their own separate comprehensive health care systems and thus do not have SHCs. Another 33 institutions did not have a SHC website as of November 20, 2019 (File S2 File.). For the remaining 549 schools, the following fields were retrieved: University name, address, website, university type, selectivity, and student demographics (i.e, percent white, African American, Hispanic and Asian).

#### 3.1.1 Student Health Center (SHC) website identification

Since the goal of this study is to analyze the reproductive health and contraceptive information provided on SHC websites, the first task we undertook was to find the SHC website, more specifically, the web address (URL) of the SHC website for a given university. We have designed and developed an algorithmic approach for this task that consists of four simple steps:

1. Construct a search query by joining the given university name with the phrase “student health center” (e.g. “Texas A & M University Central Texas student health center”).
2. Run the search query using a commercial search engine (e.g. Google Custom Search API^2^).
3. Retrieve the first result, specifically, the URL of the first result. If the URL is not from “.edu” domain, then retrieve the next URL. Repeat this until the third URL is processed. If none of the top three URLs are from “.edu” domain, then conclude that the SHC website cannot be found for this university and end.
4. Sanitize the retrieved URL to obtain the definitive URL for the SHC homepage. To do that:

a. Check if the URL redirects to another URL. If yes, then use the new URL.
b. Remove sub-URLs such as “/contacts”, “/appointments”, “/location” from the URL.

This multi-step approach was needed because there are no set standards for where and how the information related to student health services is hosted. There are no naming conventions for the SHC web addresses and as a result the SHC web addresses demonstrate high variability. For example, the web addresses for SHCs at the following five CSUs use five different naming styles:

1. California Polytechnic State University, San Luis Obispo https://hcs.calpoly.edu
2. California State University, Bakersfield https://www.csub.edu/healthcenter
3. California State University, Stanislaus https://www.csustan.edu/health-center
4. California State University, San Bernardino https://www.csusb.edu/student-health-center
5. California State Polytechnic University, Pomona https://www.cpp.edu/~health

#### 3.1.2 Assessing LARC information on SHC website

Once the SHC website is identified for a university, the LARC information provided on the SHC website is studied next. To guide this study, a rubric was developed by a certified nurse-midwife (one of the authors) and a nursing student research assistant. Two key aspects of the LARC information on SHC websites – accessibility and quality – were chosen for the rubric. These aspects of LARC information can empower college students to pursue informed decisions about LARC, ultimately making a decision with the support of a qualified healthcare provider. Accessibility of any information directly impacts its use and application. In case of SHC websites, information that is posted on their homepage has much higher accessibility than information on a web-page that is several clicks away from the SHC homepage. The quality of the provided information also impacts its use and application. In the context of this study, quality is defined as understandability of the information. If the LARC information provided on a SHC website uses simple, easy to understand language then it is more likely to be understood and in turn used. In contrast, if the LARC information on SHC website uses specialized medical terminology, then an average college student is unlikely to find it useful. There is an extensive body of research in the context of doctor-patient communication that transfers over this study [28–31]. The details of how these two aspects were quantified for a given SHC website are provided in the next two subsections.

##### LARC Information Accessibility Metric: #Clicks

The #Clicks metric was developed to answers the question: “How quickly can LARC information be reached from the SHC homepage?”. #Clicks metric captures the minimum number of clicks starting from the SHC homepage required to reach LARC content. We have designed and developed an algorithmic solution to compute this metric for any given SHC website, so as to facilitate large-scale analysis using this metric. The pseudo-code for our approach is given in Algorithm 1. At a high-level, the algorithm is designed to automate the website navigation and LARC information search process starting from a given SHC homepage. In order to find the LARC content that is closest to the SHC homepage (minimum number of clicks), this exploration is conducted in a *breadth-first* manner where all the web-pages at the same level/depth are explored before web-pages at deeper levels. To operationalize this logic, a *queue* is used to prioritize the web-pages (URLs) that have to be explored iteratively.

**Algorithm 1:**
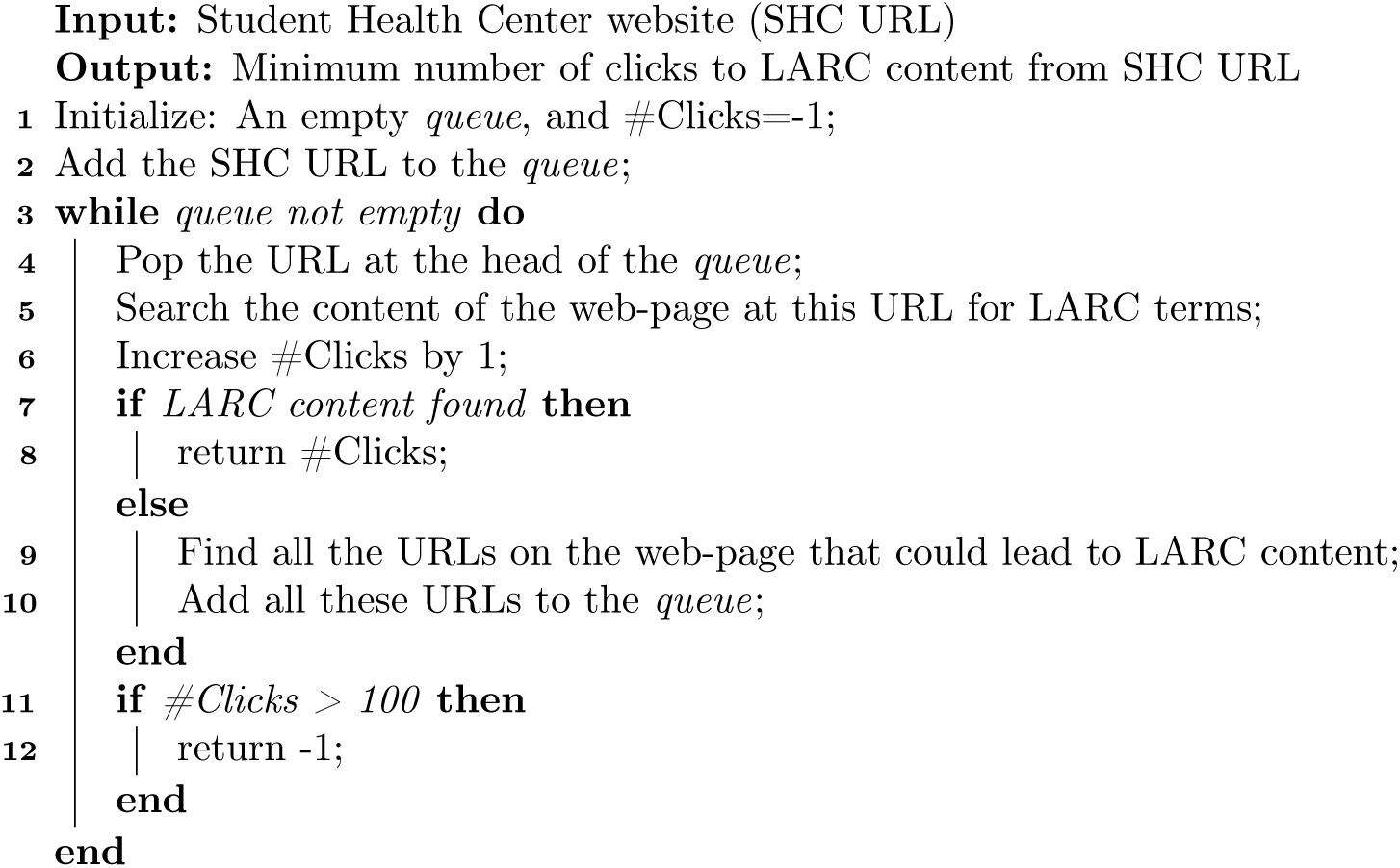
Algorithm to compute #Clicks Metric.

As is shown in Algorithm 1, the expected input to this approach is the SHC website. The URL of the SHC website is the first URL that is added to the queue (Line 2). At each iteration, the URL at the head of the queue is obtained (Line 4), and the web-page content at this URL is searched for LARC terms (Line 5). These LARC terms and phrases used in Study 1 were reused here. Every time a URL is popped from the queue and searched, the #Clicks is increased by one (Line 6) because these steps emulate the action of user clicking a link and exploring the new web-page for LARC content. The algorithm terminates as soon as the first instance of any of the LARC terms is found on a web-page (Line 8). For instance, if SHC homepage has LARC content, then the algorithm stops right after exploring the first URL (SHC homepage), and returns value 0 for #Clicks since no clicks were needed to reach LARC content. When a web-page being explored does not contain LARC content, URLs on that web-page that may lead to LARC content are identified and added to the queue (Line 9 and 10). To identify such URLs, the text of the hyperlinks on the web-page is leveraged. Specifically, if the hyperlink text contains any of the predefined keywords then the corresponding URL is added to the queue. The set of keywords was defined based on the observed trends such as SHC websites tend to provide LARC information under sections titled ‘Clinical Service’, ‘Women’s health’, ‘Reproductive health’. The program is terminated if LARC content has not been located even after exploring 100 URLs. It is assumed that LARC content is not present on this SHC website, and a special value of −1 is returned for #Clicks metric to indicate the same. The accuracy of this algorithmic approach was also analyzed and found to be 91% (Details in File S3 File.).

##### LARC Information Quality Metric: Readability

To quantify the intuition of information understandability we employ the Flesch-Kincaid readability tests^3^. The Flesch-Kincaid readability tests consist of two metrics that use linguistic properties of the textual content to estimate its readability – the Flesch Reading Ease (FRE) metric, and the Flesch-Kincaid Grade Level (FKGL) metric (formulations are given below). A higher score for FRE metric indicates easy to read material, and lower score indicates that the material is difficult to read. (Information on interpretation of FRE scores in File S4 File.) The score computed by Flesch-Kincaid Grade Level metric corresponds to U.S. grade levels. We apply both these metrics to assess the quality (understandability) of the LARC information provided on SHC websites. To operationalize this efficiently we have designed and developed computational approach that consists of four steps:

1. Construct a disjunctive query with all the LARC keywords.
2. Use a commercial search engine (e.g. Google Custom Search API) to conduct a site-specific search with the above query against the SHC website. (Site-specific search returns only those result web-pages that are hosted under the specified site, in our case, the SHC website.) Download all the results web-pages.
3. For each web-page, locate every sentence containing any of the LARC keywords, extract this sentence and 5 sentences before and after it. This step ensures that only that content which is about LARC and in the vicinity of LARC is used to compute the readability scores.
4. Count the number of syllables, words, and sentences for the content extracted in the above step^4^. Compute the Flesch Reading Ease metric:

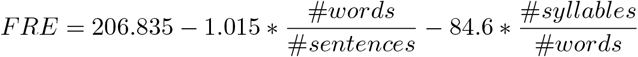 Compute the Flesch-Kincaid Grade Level metric:

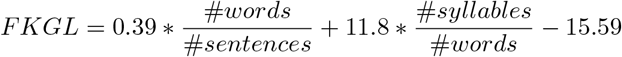

### 3.2 Study 2: Results

#### 3.2.1 SHC website identification

The seemingly simple task of finding SHC homepage for a university illustrates why many of these tasks are challenging to accomplish programmatically. The lack of standardization leads to large variance in how the SHC websites are named and organized. Many universities offer *Wellness Center* that are separate from SHC but are hard for search engines to distinguish. The SHC website identification approach described in Section 3.1.1 is designed to handle most of the above situations. To evaluate the effectiveness of this approach, the ground truth data, that is, the correct SHC websites, were manually identified for all 549 universities. When compared to this ground truth data, the SHC website identification approach is 96.36% accurate (529 out of 549 universities). Error analysis shows that majority of the identification errors are caused due to one or more of the following reasons.

1. SHC and Wellness Center: In case of some universities the Wellness Center website also contains many of the search query terms and is ranked higher than SHC website by the search engine. For instance, the search query “Alabama A&M University student health center” retrieves their Wellness Center website at top rank and the SHC website is ranked second. The reason for this is that the Wellness Center homepage mentions “student health” several times including in its title. The SHC homepage, on the other hand, has the following title “John and Ella Byrd McCain Health and Counseling Center - Alabama A&M University” and just one instance of “student health” on the homepage.
2. Inconsistent Domain Name: Some SHC website use different domain name than university. For example, the SHC website for University of Maine is https://northernlighthealth.org/Locations/Eastern-Maine-Medical-Center/Locations/Primary-Care-Umaine. The computational approach however identifies a different URL under umaine.edu domain as the SHC website: https://umaine.edu/studentlife/parents-and-family/campus-healthcare/. This web-page provides information about the SHC but it is not the SHC website; it does not even include a hyperlink to the correct SHC website.
3. Mutable Ground Truth Data: Due to the dynamic nature of online content, the ground truth data for the task at hand may change. For instance, the SHC website for Wayne State College few months ago was https://www.wsc.edu/info/20026/campus_life/80/student_health_office but now it has changed to https://health.wayne.edu. The algorithmic approach retrieves the correct URL but it does not match the now defunct ground truth URL.

#### 3.2.2 LARC information accessibility measure: #Clicks

The accessibility measure, #Clicks, was applied to all 549 university SHC websites, to compute the minimum number of clicks needed to reach LARC information on each SHC website. The corresponding results are summarized in Table 2. A very small fraction of the universities (2.36%) place LARC information on SHC homepage. A quarter of the universities (25.15%) place LARC information 1 to 4 clicks away from the SHC homepage. Majority of the schools (72.49%) do not provide any LARC information on their SHC website. If we consider 0 – 1 clicks as high accessibility, 2 – 3 as moderate accessibility, and 4+ as low accessibility, then approximately the same number of universities (70) are in the high and moderate accessibility range. Only one university SHC is in the low accessibility range.

**Table 2.** #Clicks Metric: Minimum no. of clicks to LARC content from SHC homepage

| #Clicks | Number of universities (Percentage) |
| --- | --- |
| 0 | 13 (2.36%) |
| 1 | 64 (11.65%) |
| 2 | 60 (10.92%) |
| 3 | 13 (2.36%) |
| 4 | 1 (0.7%) |
| No LARC content | 398 (72.49%) |

#### 3.2.3 LARC information quality measure: Readability

The results for the two readability metrics, Flesch Reading Ease (FRE) and Flesch-Kincaid Grade Level (FKGL), are summarized in Table 3. Of the 549 universities, 171 university SHC websites have LARC content. Majority of this subset (75% as per FRE, and 81% as FKGL) provide LARC information in plain English (reading level below 10th grade). Only 9% as per FRE, and 5% as per FKGL, universities had information that would be considered difficult, higher than 12th grade level (File S5 File provides a list of these universities.). The overall trend is that when LARC information is provided on SHC website, it is generally readable/understandable to college students.

**Table 3.**
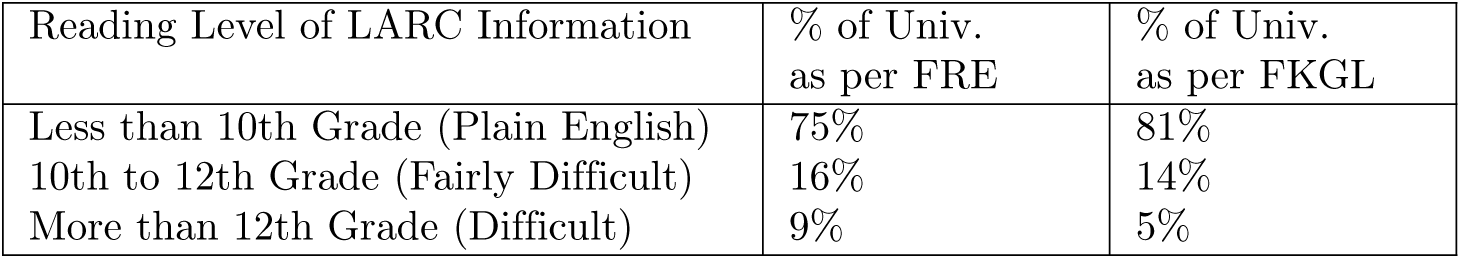
Reading Level of LARC Information on SHC website

## 4 STUDY 3: Factor analysis of LARC measures with regional and other external factors

Access to LARC information can be associated with a myriad of factors ranging from the institutional factors to contextual factors related to location [16, 17]. We examine a few institutional level demographic variables (racial composition, gender composition) and urbanicity at the county level as key factors related to university providing LARC information on SHC website. Institutional level data is from Integrated Postsecondary Education Data System and county level urbanicity is based on designations from the Center of Disease Control (https://www.cdc.gov/nchs/data_access/urban_rural.htm).

### 4.1 Study 3: Methods

Our main analyses examine the associations between institutional demographic variables and county urbanicity and having LARC information on the SHC website. We estimate the association using standard logistic regression models. We present a series of bivariate associations and multivariate association where we account for demographic and urbanicity characteristics simultaneously and control for census region. This analysis was conducted on same set of universities (549) as Study 2.

For the subset of 171 universities where LARC information was found on their SHC website, we also analyzed the associations between demographic and urbanicity factors with #Clicks (the minimum number of clicks to the LARC information on SHC website) and the readability scores of the LARC information. Given the small sample we examine difference using Chi-squared test for this analysis.

### 4.2 Study 3: Results

Table 4 presents odds ratios and 95% confidence intervals for the association between university demographic and location characteristics and having LARC information on the SHC website. Columns 1-4 present bivariate relationship and column 5 presents the multivariate relationship. Results from column 1 and 3 suggests that universities having high rates of the student body who are African American (AA) or female are less likely to provide LARC information on their SHC website. Results from column 4 suggest that universities situated in more rural settings are less likely to provide LARC information on their SHC websites. These three characteristics remain statistically significant in multivariate regressions though the two demographic characteristics become only marginally significant (column 5).

For the subset of universities that provide LARC information on their SHC websites, we examine #Clicks values and readability scores. LARC information was usually found between 0 to 4 clicks. Most schools with LARC information have details within 1 click from the SHC homepage. There were limited systematic differences, except that at universities with the highest proportion of African American students the number of clicks was lower (*χ*^2^(6, *N* = 148) = 15.45, *p* < 0.05). There were no systematic differences for the readability scores.

**Table 4.**
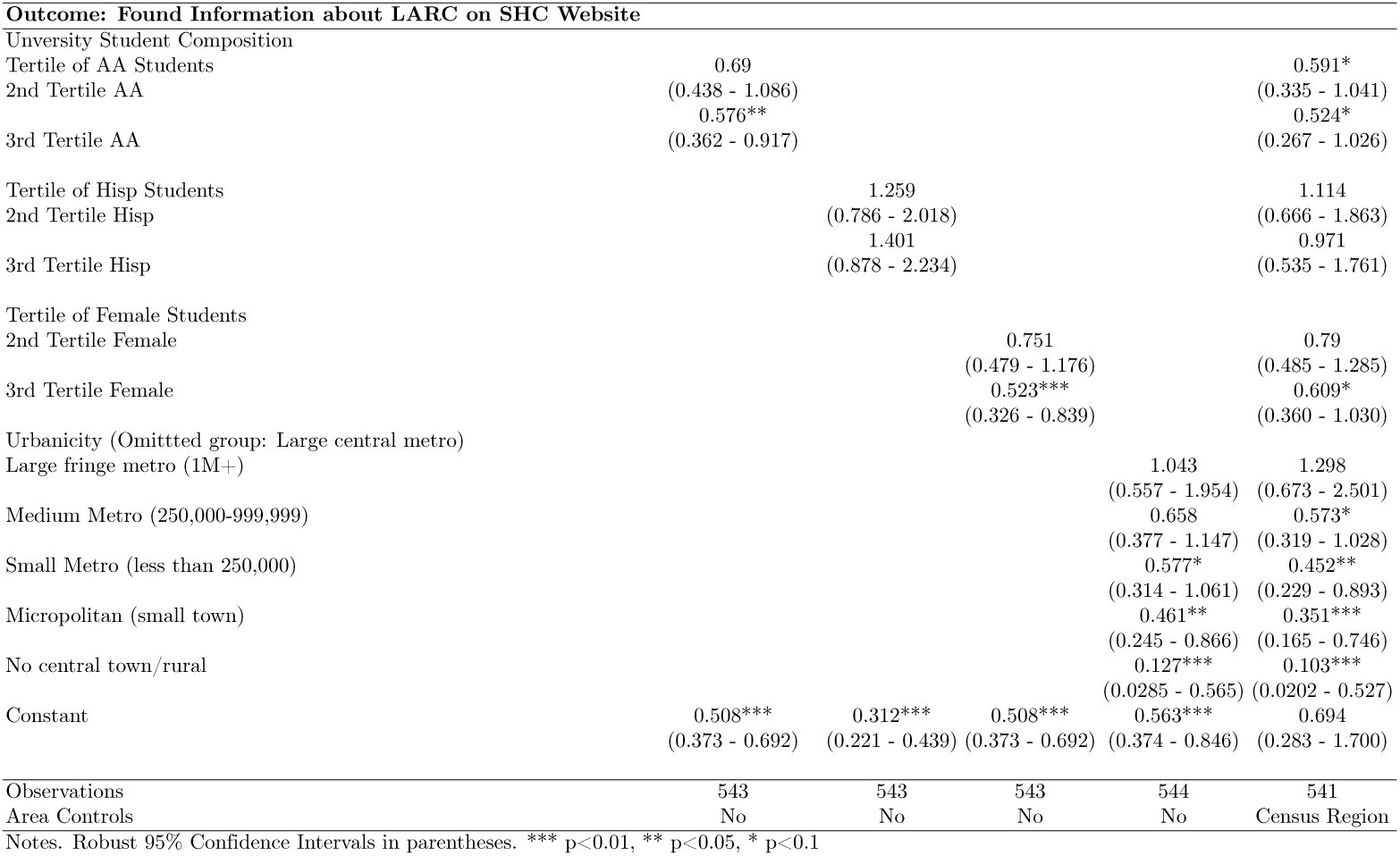
University Characteristics Associated with Having LARC information on SHC Website

## 5 Discussion

Together the results of our three studies suggest that LARC methods, despite being among the most effective forms of contraception, are less well represented on SHC websites compared to other common reproductive health services such as EC, condoms, and Pap smears. In addition to finding that LARC methods were mentioned less often than the other methods, Study 1 findings suggest that manual annotation may not be the gold standard for research questions regarding online health information access. Study 2 shows that while only 27.5% of the SHCs at U.S. public universities mention LARC methods on their websites, at SHCs where LARC information is provided it has fair accessibility in terms of navigation and readability. In addition, in Study 3 we found that public universities with higher proportions of African American students and female students were less likely to mention LARC information on their SHC websites. Our results suggest that institutional demographic characteristics are associated with access to LARC information for college students. Most importantly, informational disparities may hinder access to and use of LARC methods in ways that could potentially exacerbate pre-existing disparities by putting students at increased risk for unplanned pregnancy.

Systematic manual and automated reviews as we have done here may offer an innovative alternative to surveys in the assessment of reproductive health information provided by SHCs. When previous studies have relied on surveys, there has been the risk of low response rates and likely highly select responses from universities with strong SHCs. In addition, surveys of staff and students at SHCs may not accurately reflect the experiences of the full range of students attempting to access contraceptive information through the SHCs. Our results suggest that human annotation is variable even with simple predetermined rubrics, and thus automated methods may promise greater reliability, especially for online information. The above studies demonstrate that the data gathering process for online LARC information can be automated with high accuracy. Study 3 results are supported by literature showing that African American women access prescription contraceptive methods such as LARC at lower rates than white women even after passage of the ACA [3]. There is also evidence that care provided at rural public health clinics may pose unnecessary barriers to LARC access or simply not provide LARC services [32].

### 5.1 Strengths and limitations

This study has several important strengths, the first being that it applies an interdisciplinary approach to a traditional public health question, leveraging the strengths of several disparate fields for a new and innovative approach. Moreover, our results show that automated approaches can be used to efficiently scale data gathering efforts in a systematic way, which can support longitudinal tracking studies that observe the changes in online informational health data provided by organizations. Specifically, while the results from the SHC website identification task revealed a lack of standardization for SHC web addresses, the now-vetted algorithm could be deployed periodically in the future to capture and monitor the dynamic nature of online LARC information. Furthermore, we have shown that computational approaches could be easily generalized to gather data from SHC websites about other health topics for example the prevention of sexually transmitted infections such as HIV.

This study also has several limitations. Firstly, there is the contextual issue of ACA coverage, which mandates contraceptive provision by SHCs but does not require that all types of contraceptives be available and covered at SHCs. An inherent assumption of our study is that one could expect accurate, accessible information about LARC methods to be offered on an SHC website even if the services are not provided by the SHC. The rationale for this is that posting the information on the website has a positive impact on choice and safety for students [4] and is essentially low-cost or free. Yet given the information available on the websites (or lack thereof), it was not possible for us to fully ascertain which schools may be providing these services in their clinics. Additional research is needed to understand the link between LARC information offered on SHC website and service provision in SHC clinics. Another key limitation is that only a few variables were examined as factors associated with LARC information and the evaluation occurred at one point in time. To understand whether these associations are casual, future studies would be needed in order to build a panel of relevant changes in student demographics and LARC information.

## 6 Conclusion

SHCs have the potential to positively impact health-related outcomes for U.S. college students by providing access to high-quality contraceptive information on their websites, including information on LARC. Improving the low rate at which SHC websites currently provide LARC information would be an effective and low-cost method of improving health outcomes and well-being for college students. In addition, interdisciplinary automated approaches for data collection such as the one we have developed here hold promise for the future study of public health questions involving online information. With repeated use, an automated approach could provide a reliable method for monitoring changes in online health-related information over time. In contrast to traditional survey research, our systematic approach revealed that even groups perceived to be socioeconomically privileged, such as U.S. college students, can experience different levels of access to basic health information and services such as LARC methods based on predictors such as race and urban or rural geographic area. This hidden disparity is often overlooked based on the commonly held assumption that the provision of online information related to a health service can be interpreted as evidence of the provision of that service, without taking into account the accessibility of the online information regarding services for the target population. Future research is needed to test that assumption and explore the role of online information provision and accessibility in health service access, particularly for under-served populations.

## Supporting information

**S1 File. Table S1.** List of 200 universities sampled for manual annotation in Study 1.

**S2 File. Table S2.** List of 33 universities that did not have SHC website as of Nov 20, 2019.

**S3 File. File S3.** Details about evaluation of #Clicks algorithm.

**S4 File. Table S4.** Flesch Readability Ease (FRE) score interpretation table.

**S5 File. Table S5.** List of Universities with Reading Level higher than 12th grade for LARC information on their SHC website.

## Acknowledgments

We thank students who contributed to data collection [list of students in alphabetical order]: Sabrina Gonzaga, Fungai Gora, Byron Mills. Student time was supported by a training grant from the National Institutes of Health, R25 MD011714.

## Footnotes

1 If instead of using 0.5 as an uncertainty value in the rows with a disagreement we simply treat them as missing values, the overall statistical t-test is t(156) = −1.84, *p*<.05, one-tailed.

2 https://developers.google.com/custom-search/v1/overview

3 https://en.wikipedia.org/wiki/Flesch-Kincaid_readability_tests

4 NLP library was used for this: spaCy, https://spacy.io/

